# A revised understanding of *Tribolium* morphogenesis further reconciles short and long germ development

**DOI:** 10.1101/232751

**Authors:** Matthew A. Benton

## Abstract

In *Drosophila melanogaster*, the germband forms directly on the egg surface and solely consists of embryonic tissue. In contrast, most insect embryos undergo a complicated set of tissue rearrangements to generate a condensed, multi-layered germband. The ventral side of the germband is embryonic, while the dorsal side is thought to be an extraembryonic tissue called the amnion. While this tissue organisation has been accepted for decades, and has been widely reported in insects, its accuracy has not been directly tested in any species. Using live cell tracking and differential cell labelling in the short germ beetle *Tribolium castaneum*, I show that most of the cells previously thought to be amnion actually give rise to large parts of the embryo. This process occurs via the dorsal-to-ventral flow of cells and contributes to germband extension. In addition, I show that true ‘amnion’ cells in *Tribolium* originate from a small region of the blastoderm. Together, my findings show that development in the short germ embryos of *Tribolium* and the long germ embryos of *Drosophila* is more similar than previously proposed. Dorsal-to-ventral cell flow also occurs in *Drosophila* during germband extension, and I argue that the flow is driven by a conserved set of underlying morphogenetic events in both species. Furthermore, the revised *Tribolium* fatemap that I present is far more similar to that of *Drosophila* than the classic *Tribolium* fatemap. Lastly, my findings show that there is no qualitative difference between the tissue structure of the cellularised blastoderm and the short/intermediate germ germband. As such, the same tissue patterning mechanisms could function continuously throughout the cellularised blastoderm and germband stages, and easily shift between them over evolutionary time.

**Author summary:** In many animals, certain groups of cells in the embryo do not directly contribute to adult structures. Instead, these cells generate so-called ‘extra-embryonic tissues’ that support and facilitate development, but degenerate prior to birth/hatching. In most insect species, embryos are described as having two major extra-embryonic tissues; the serosa, which encapsulates the entire embryo and yolk, and the amnion, which covers one side of the embryo. This tissue structure has been widely reported for over a century, but detailed studies on the amnion are lacking. Working in the beetle Tribolium castaneum, I used long-term fluorescent live imaging, cell tracking and differential cell labelling to investigate amnion development. In contrast to our current understanding, I show that most cells previously thought to be amnion actually form large parts of the embryo. In addition, I show how these cells ‘flow’ as a whole tissue and contribute to elongation of the embryo, and how only a relatively small number of cells form the actual amnion. Lastly, I describe how my findings show that despite exhibiting substantial differences in overall structure, embryos of Tribolium and the fruit fly, Drosophila melanogaster, utilise a conserved set of morphogenetic processes.

## Introduction

Insects are the most speciose phylum of animals and display remarkable diversity in adult morphology [1]. Insect embryo development is also very diverse, particularly in the stages leading to the formation of the elongated, segmented embryo (called the germband) [2]. The molecular and morphogenetic basis of this process is best understood in the fly *Drosophila melanogaster*. In this species, a predominantly hierarchical chain of patterning events specifies nearly all segments more-or-less simultaneously at the syncytial blastoderm stage [3]. Cellularisation takes place near the end of this process, after which point morphogenetic events such as germband extension (GBE) occur (see Fig 1 for schematic summary). The *Drosophila* mode of development is termed long germ development and is fairly representative of most true flies [4]. In contrast, the vast majority of insects undergo short or intermediate germ development, meaning that only a handful of segments are specified at the blastoderm stage and the remaining segments are specified sequentially as the germband elongates [5].

**Fig 1.**
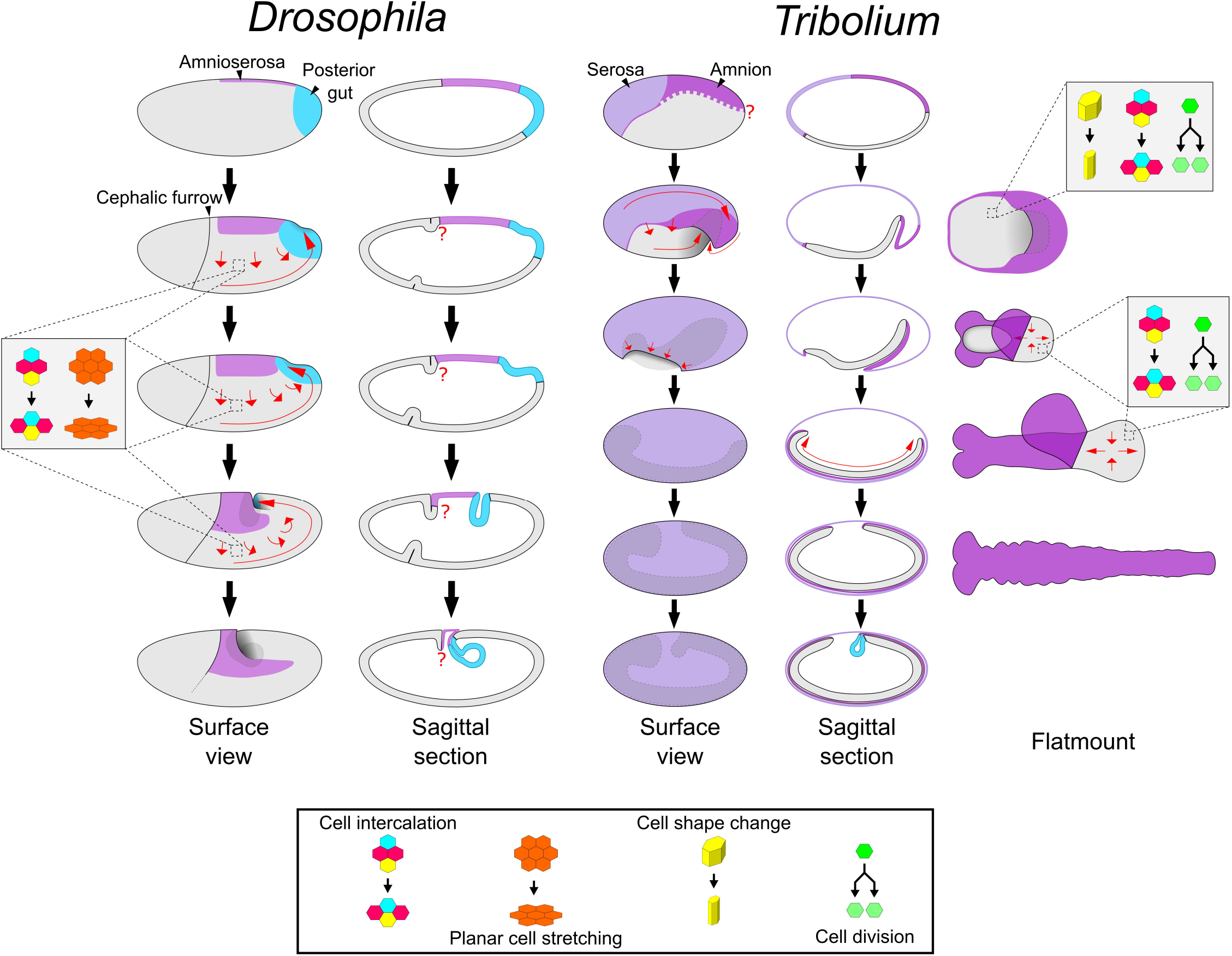
Schematics of development in *Drosophila* and *Tribolium*. The two left columns show schematics of *Drosophila* embryos from the uniform blastoderm stage to the extended germband stage. The right three columns show schematics of *Tribolium* embryos at comparable developmental stages. The schematics in the right-most column depict dissected, flatmounted embryos. Red arrows display cell/tissue movement. The question marks highlight two regions (the *Drosophila* embryo/amnioserosa border in the cephalic furrow region, and the dorsoventral position of the *Tribolium* embryo/amnion border) where the tissue boundaries are unknown/undescribed. Several features have been omitted, including the yolk, mesoderm gastrulation, anterior gut formation and appendage formation. The *Drosophila* fatemap is based on data from [18] and the references therein. Refer to text for additional details.

Short germ development has been best studied in the beetle *Tribolium castaneum*, and recent research has shown that development in this species is more similar to *Drosophila* than previously thought. In *Drosophila*, GBE is predominantly driven by the mediolateral intercalation of ectodermal cells (i.e. convergent extension), although cell deformation along the anterior-posterior (AP) axis and cell divisions are also involved [6–11]. In contrast to this, *Tribolium* germband elongation was previously thought to be driven by the so-called ‘growth zone’ at the posterior of the germband [12]. Now, however, it is clear that *Tribolium* germband elongation is also predominantly driven by mediolateral cell intercalation (see Fig 1 for schematic summary of *Tribolium* development) [13–15]. Furthermore, in both *Tribolium* and *Drosophila*, this intercalation requires the striped expression of a specific group of Toll genes (so-called Long Toll/Loto class genes) [16,17].

It is highly likely that germband elongation mediated by cell intercalation is homologous in these two species, and probably in other arthropods, as well [17]. As such, I will hereafter refer to *Tribolium* ‘germband elongation’ as ‘germband extension’/GBE, unifying the *Drosophila /Tribolium* terminology. In addition, as there is no evidence for a qualitatively different ‘growth zone’ in *Tribolium* (i.e. a specialised zone of volumetric growth), I will refer to the posterior unsegmented region as the ‘segment addition zone’ (SAZ) [19–21].

Despite the similarities described above, there are substantial differences in the embryonic fatemaps of these two species (Fig 1). In *Drosophila*, almost the entire blastoderm is fated as embryonic tissue, and only a small dorsal region is fated as extraembryonic tissue (termed the amnioserosa) [18]. In contrast, in *Tribolium*, roughly the anterior third of the blastoderm gives rise to an extraembryonic tissue called the serosa [22]. Of the remaining blastoderm, a large dorsal region is thought to give rise to a second extraembryonic tissue called the amnion, with only the remaining ventral tissue giving rise to the embryo itself [23–25]. Like the amnioserosa, the serosa and the amnion are proposed to support the embryo during development, but are thought to degenerate prior to hatching and not contribute to any larval or adult structures [19,26,27].

*Drosophila* and *Tribolium* also exhibit dramatic differences in the morphogenetic events occurring during early development (Fig 1). When GBE occurs in *Drosophila*, the germband stays at the surface of the egg and the amnioserosa largely remains in place. In *Tribolium*, on the other hand, germband extension begins with a process called embryo condensation, during which the embryonic ectoderm and presumptive amnion (together termed the ‘germ rudiment’) form the germband (see Fig 1; for a detailed description see [14,28]). Several concurrent morphogenetic events underlie embryo condensation. The embryonic ectoderm condenses towards the ventral side of the egg via both mediolateral cell intercalation and a cuboidal-to-columnar cell shape transition. Simultaneously, epithelial folding and tissue involution occurs, causing the presumptive amnion to fold over the embryonic ectoderm. During these movements, the serosa cells undergo a cuboidal-to-squamous transition to spread over the entire egg surface. The final stage of embryo condensation coincides with closure of the serosa (serosa window stage), which appears to involve a supracellular actomyosin cable [14].

The differences in fatemap and tissue folding described above show that both fatemap shifts and reductions in early morphogenetic events have contributed to the evolution of the long germ mode of development found in *Drosophila*. However, it is important to note that *Drosophila*, regarding the extraembryonic tissues, represents an extreme case of reductive evolution, which is characteristic only for higher cylorrhaphan flies [29]. More basally branching flies form both an amnion and a serosa, while still exhibiting the long germ mode of development (for a review see [26]). For example, in the scuttle fly *Megaselia abdita*, both an amnion and serosa form, but while the serosa spreads over the egg surface as in *Tribolium*, the amnion remains at the dorsal side of the embryo, similar to the *Drosophila* amnioserosa [30–33]. Such intermediate topologies help to explain the evolution of the situation in *Drosophila*, where all extraembryonic cells remain at the dorsal side.

Understanding how these differences evolved is integral to understanding the short-to-long germ transition, but in order to study how this occurred, we first need to understand how these tissues develop in each species. The form and function of the *Tribolium* serosa has been analysed in several studies [22,34,35]. The amnion, on the other hand, has proven harder to analyse, and the precise embryo/amnion boundary at the blastoderm stage is unknown. However, a defined boundary between embryo and amnion has been proposed to exist from when the germband forms (Fig 1) [23]. Cells in the ventral half of the germband (ventral with respect to the germband dorsoventral [DV] polarity, but dorsal with respect to the egg) are thought to give rise to all embryonic structures, while cells in the dorsal half of the germband (dorsal with respect to the germband DV polarity, but ventral with respect to the egg) are thought to form the amnion [25,36,37]. This germband structure has been described in many insects over the past century and is proposed to represent the core conserved structure of short/intermediate germ embryos (reviewed in [2,38,39]). However, the proposed boundary between cells fated to become embryo and those fated to become amnion has not been directly tested.

Here, I investigate the development of the presumptive amnion in *Tribolium* using a combination of fluorescent live imaging and fate mapping techniques. To my great surprise, I find that the majority of the cells previously described as ‘amnion’ actually form large parts of the embryo proper. Using fate-mapping experiments, I show that true ‘amnion’ cells originate from a very small domain of the blastoderm, just as the *Drosophila* amnioserosa cells do. I also show that the movement of cells from the ‘amnion’ side of the germband to the ‘embryo’ side of the germband occurs via the large scale flow of the ectodermal epithelium. Lastly, I describe the underlying causes of this flow, and show how this tissue movement is likely homologous to the dorsal-to-ventral tissue flow that occurs during *Drosophila* GBE.

## Results

### Live cell tracking reveals movement of ‘amnion’ cells into the embryo

To examine the development of the *Tribolium* presumptive amnion in detail, I carried out high resolution live imaging of embryos transiently labelled [14] with a fluorescent histone marker (H2B-venus) to label nuclei. My goal was to track presumptive amnion cells from the blastoderm stage onwards. However, it was not possible to accurately track the majority of cells throughout embryo condensation and GBE, due to the extensive morphogenetic rearrangements that take place during this process. Instead, I focused on the stage immediately following condensation when the germband has formed, and analysed the embryonic region where the presumptive amnion is closest to the surface of the egg. Specifically, I tracked over 200 presumptive amnion cells from the central region of the germband from the closure of the serosa window until after the formation of the thoracic segments (over 11 hours of development; Fig 2 and S2 Movie). As previously described [14], the germband and yolk exhibit pulsatile movements during this period, as well as rotating within the serosa (S1 Movie).

**Fig 2.**
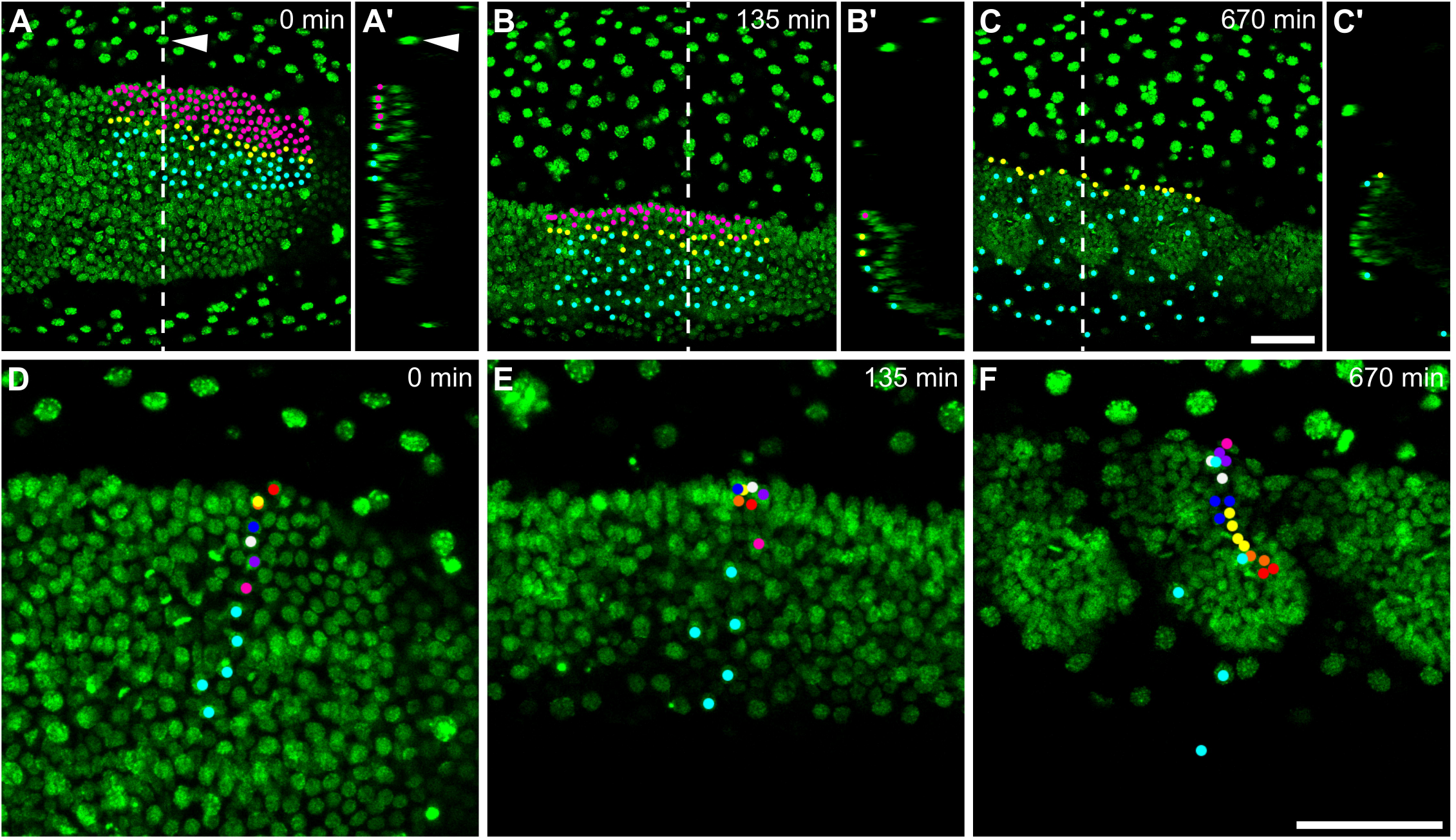
Live cell tracking reveals contribution of ‘amnion’ cells to embryonic tissue. A-F) Time series from fluorescent live imaging of a *Tribolium* embryo expressing H2B-venus. The serosa nuclei located above the germband have been manually removed from these frames (by deleting them from individual z-stack slices), but left in the surrounding territory (arrowhead in (A+A’)). (A’-C’) show optical transverse sections of the respective frame at the position shown by the dashed line (the surface of the egg is to the left). In (A-C), all nuclei that lie in a region of the ‘amnion territory’ in (A) have been tracked and differentially labelled depending on whether they become part of the embryo (magenta; labels disappear when nuclei join the germband), become located at the edge of the germband (yellow) or remain in the ‘amnion territory’ (cyan). In (D-F), a line of nuclei that lie in the ‘amnion territory’ in (D) have been tracked and differentially labelled depending on whether they become part of the embryo (coloured points; daughter cells are labelled in same colour as parent) or remain in the ‘amnion territory’ (cyan; no division takes place). Note that in panel (D), the orange spot is mostly hidden below the yellow spot because the nuclei in that region are partially overlapping when viewed as projections. The first frame of the timelapse was defined as timepoint 0. In (A-F), embryos are oriented based on the AP/DV polarity of the egg with anterior to the left and dorsal to the top. (A-C) are maximum intensity projections of one egg hemisphere. (D-F) are average intensity projections of 46 microns to specifically show the germband. Scale bars are 50 μm.

The presumptive amnion initially consists of many tightly packed cells, which become increasingly spread out during GBE (S2 Movie, Fig 2(A-C)). However, rather than remaining restricted to the ‘amnion territory’, many of the tracked cells moved around the edge of the germband into the ‘embryo territory’. Differential labelling of tracked cells clearly showed that these cells that moved around the germband edge became part of the embryo proper (S2 Movie and Fig 2(A-C)). The cells that joined the ‘embryo territory’ became tightly packed, continued to divide, and formed embryonic structures (S3 Movie and Fig 2(D-F)). In contrast, cells that remained in the ‘amnion territory’ became squamous and stopped dividing. The nuclei of these latter cells became enlarged (S3 Movie and Fig 2(D-F)), suggesting that they underwent endoreplication to become polyploid, as seen in the *Tribolium* serosa and in the *Drosophila* amnioserosa [18,24]. In addition, several germband nuclei underwent apoptosis (S3 Movie) as has been described in fixed embryos [40]. These results show that many of the cells previously thought to constitute extraembryonic amnion give rise to embryonic structures.

Since the epithelium formerly termed ‘amnion’ is made up cells that will variously form amnion, dorsal ectoderm and dorsolateral ectoderm, it is not accurate for the entire tissue to be called ‘amnion’. Therefore, I will refer to this part of the germband as the ‘dorsal epithelium’, based on the tissue’s location at the dorsal side of the germband (with respect to the DV polarity of the germband rather than the egg). This term ‘dorsal epithelium’ is simply a spatial designation, and comes with no implicit assumptions about the identity of the tissue nor the final fate of the tissue. It is also important to keep in mind that the dorsal epithelium is continuous with the ventral epithelium.

### Differential cell labelling confirms widespread dorsal-to-ventral cell movement

My next question was whether the movement of cells from the dorsal epithelium to the ventral epithelium occurs throughout the AP axis or is just limited to the thoracic region. The extensive movements of the germband made it difficult to track individual cells accurately at the anterior and posterior poles. To overcome this problem, I combined differential cell labelling with long term fluorescent live imaging to follow small groups of nuclei throughout development. Specifically, I microinjected mRNA encoding a nuclear-localised photoconvertable fluorescent protein (NLS-tdEos) into pre-blastoderm embryos to uniformly label all nuclei, then photoconverted a small patch of nuclei at different positions along the AP axis at the final uniform blastoderm stage. I then performed long term confocal live imaging of both the unconverted and photoconverted forms of the fluorescent protein throughout the period of GBE (or longer). Unlike that of *Drosophila*, the *Tribolium* egg shell does not show any dorsoventral (DV) polarity, and I was therefore unable to specifically target particular locations along the DV axis. Instead, I opted for a brute-force approach and performed the photoconversion experiment at unknown DV positions for 50-150 embryos at each of the following AP positions: 75% egg length (EL) from the posterior pole, 50% EL, 25% EL, and close to the posterior pole. I then used the resulting live imaging data to determine the approximate DV position of the photoconverted cells. Using a new live imaging set up (see Materials and Methods), I obtained the same range of hatching rates as I typically obtain for other microinjection experiments (approximately 80%, [14]), even after continuous confocal live imaging for almost the entirety of *Tribolium* embryonic development (3.5 days; S4 Movie). Both unconverted and photoconverted protein persisted throughout germband extension and retraction, although fluorescent signal faded over time. I have included various examples from this data set in S1-S3 Figures. In addition, I have made the raw confocal data for a large number of timelapses available online (>300 embryos, >700 GB of data [41]) for the benefit of the community. This data will likely prove valuable for a wide range of research projects.

When I examined clones initially located in the dorsal epithelium, I found that movement of cells from the dorsal epithelium to the ventral epithelium occurred throughout the posterior of the embryo during GBE (Fig 3(A-F), S5 Movie). I also observed the same movements at the anterior of the germband (Fig 3(G-J)), although I have focused my analysis on the middle and posterior parts of the embryo. Together with the cell tracking data, these results show that most of what was previously thought to be ‘amnion’ is in fact embryonic tissue, and that cells move from the dorsal epithelium to the ventral epithelium throughout the germband (summarised in Fig 3(K,L)).

**Fig 3.**
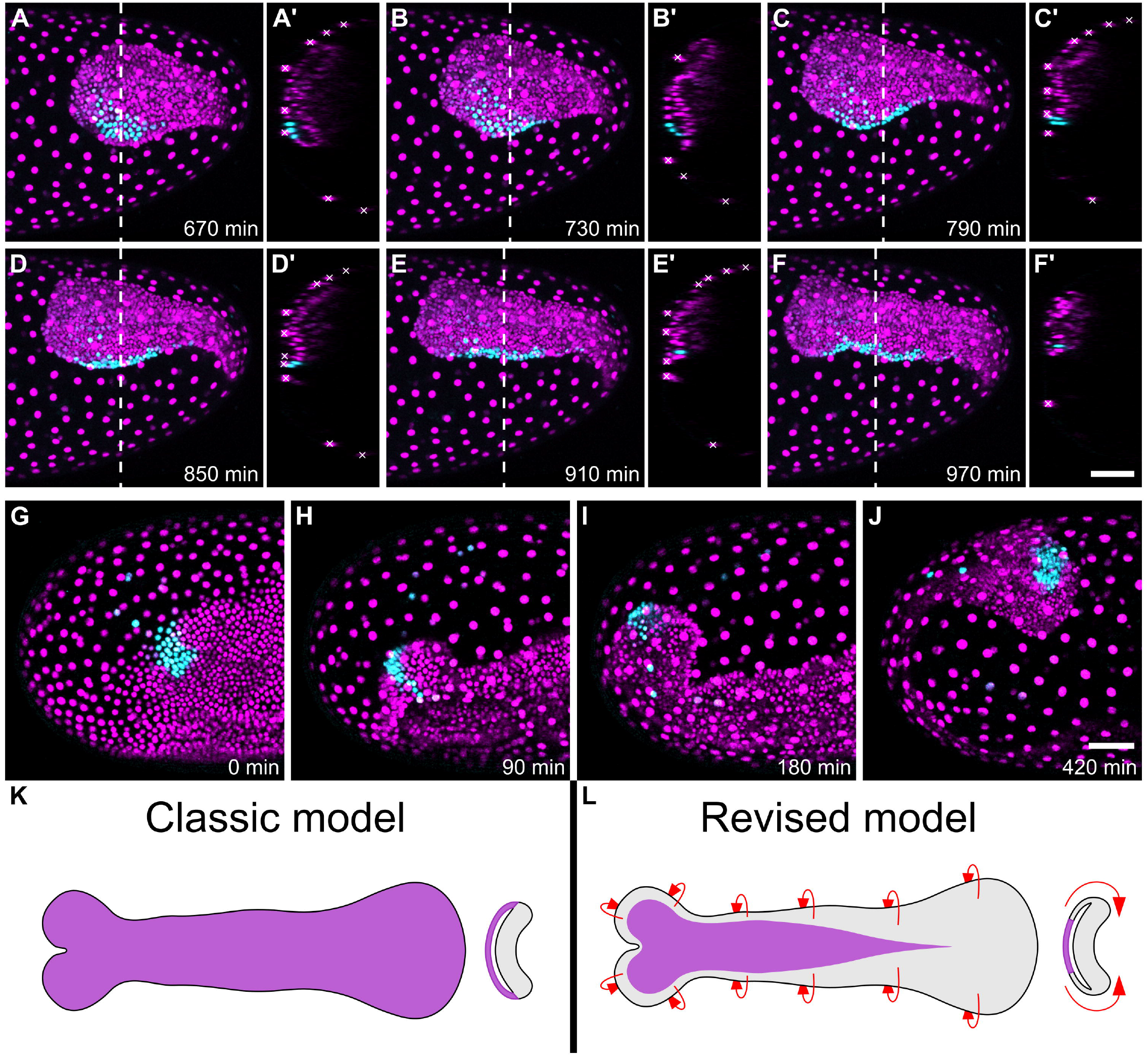
Differential cell labelling reveals widespread movement of cells from the dorsal epithelium to the ventral epithelium. (A-J) Time series from fluorescent live imaging of two *Tribolium* embryos expressing NLS-tdEos showing unconverted protein (magenta) and photoconverted protein (cyan). In (A-F’) a patch of nuclei at the posterior-dorsal region of the blastoderm were photoconverted. Panels (A-F) show the posterior region of the germband during late GBE and panels (A’-F’) show optical transverse sections made at the position of the dashed line at each timepoint (roughly following the same nuclei). Serosa nuclei are marked by white crosses in the transverse sections. In (G-J), a patch of nuclei at the anterior-lateral region of the blastoderm were photoconverted. Panels (G-J) show the anterior of the germband during condensation and GBE. In both embryos, all converted nuclei are initially located in the dorsal epithelium, but most move into the ventral epithelium during GBE. (K-L) Schematics showing the classic and revised models of the *Tribolium* germband (presumptive amnion is shown in purple, presumptive embryo is shown in grey, red arrows show the newly discovered tissue flow). The first frame of the timelapses was defined as timepoint 0. In (A-J), embryos are oriented based on the AP/DV polarity of the egg with anterior to the left and dorsal to the top. In (A’-F’), the surface of the egg is oriented to the left. In (K-L), schematics show flatmounted germbands with the focus on the dorsal epithelium, the anterior to the left and the orthogonal sections are oriented with the dorsal half of the germband to the left. (A-J) are maximum intensity projections of one egg hemisphere. Scale bars are 50 μm.

### Mediolateral cell intercalation occurs throughout GBE

During my live imaging, ectodermal cell clones became elongated along the AP axis over time, as previously reported in a *Tribolium* study that used a non-live imaging cell clone method [15]. However, this study found that “labelled ectodermal cells … rarely mix with unlabelled cells” even as clones became greatly elongated [15]. In contrast, I frequently observed non-converted nuclei in the midst of labelled nuclei (Fig 3, S1-S3 Figs).

To test whether the pattern I observed was caused by mediolateral cell intercalation, I tracked the nuclei of abutting rows of ectodermal cells in the SAZ during formation of the abdominal segments (50 cells in total, tracked for 3.5 hours; Fig 4 and S6 Movie). This analysis clearly showed that, as during embryo condensation [14]. cells intercalated between their dorsal and ventral neighbours. Together with the photoconversion dataset, these results show that extensive mediolateral cell intercalation takes place throughout GBE to drive the convergent extension of the ectoderm.

**Fig 4.**
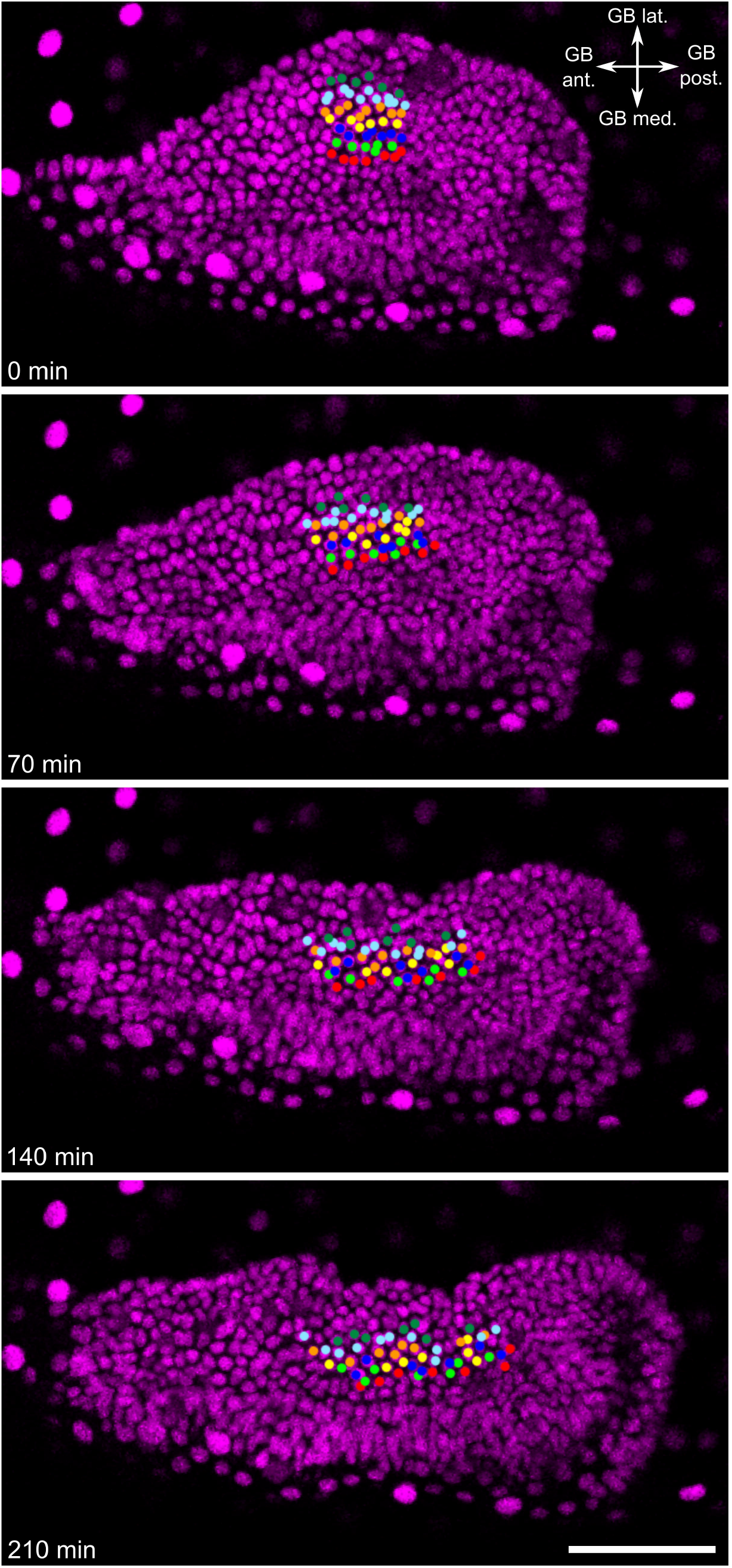
Mediolateral cell intercalation occurs in the SAZ during GBE. Time series from fluorescent live imaging of a *Tribolium embryo* expressing NLS-tdEos showing the SAZ during abdominal segment formation. Coloured points mark tracked nuclei. During the timelapse, nuclei underwent apicobasal movement but I observed no cell delamination. Note that the same embryo is shown in Fig 3(A-F). The first frame of the timelapse was defined as timepoint 0. The embryo is oriented based on polarity of the visible region of the germband. Panels show maximum intensity projections of 15 μm to specifically show the germband. Abbreviations as follows: germband (GB); anterior (ant); posterior (post); medial (med); lateral (lat). The scale bar is 50 μm.

### *Tribolium serpent* may mark true ‘amnion’

As described earlier, cells that remained in the dorsal epithelium became squamous, and this cell shape change occurred progressively along the AP axis (Fig 5(A)). This change in cell shape may be a sign of maturation of true ‘amnion’. While characterising cell fate markers, I found that the *Tribolium* ortholog of the GATA factor *serpent* (*Tc-srp*) exhibited spatial and temporal expression dynamics that were very similar to those of the potential ‘amnion’ (i.e. progressive flattening of cells, Fig 5(B), S4 Fig).

**Fig 5.**
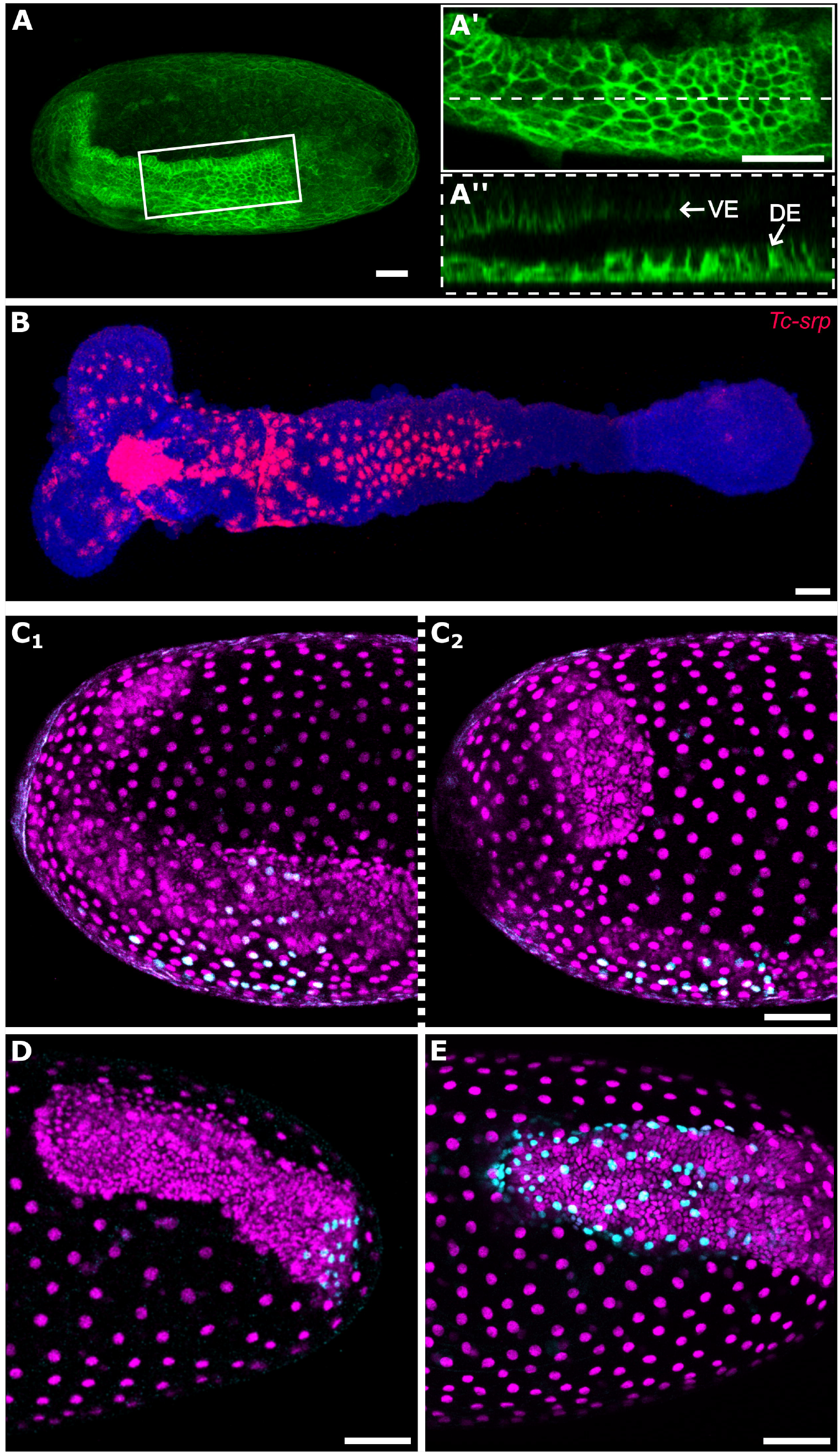
Development of the putative amnion. (A-A’’) *Tribolium* embryo transiently expressing the membrane marker GAP43-YFP. (A) shows an overview of the whole egg, (A’) shows the dorsal epithelium of the same embryo at the position of the white box, (A’’) is an optical sagittal section at the position of the dashed line in (A’) showing the apical-basal height of cells of the dorsal epithelium. (B) *Tc-srp* (red) expression in a flatmounted *Tribolium* germband also showing nuclei (DAPI, blue). The strong *Tc-srp* signal in nuclei may suggest nuclear or peri-nuclear localisation of the transcript, or it may be due to the cell body being flattened. Aside from the strong patch of anterior medial expression (which is from cells beneath the embryonic ectoderm), all visible expression is in the putative amnion epithelium. (C-E) Extended germband stage *Tribolium* embryos transiently expressing NLS-tdEos showing unconverted protein (magenta) and photoconverted protein (cyan). In each embryo, the clone of converted cells spans the entire amnion. (C_1-2_) show both sides of the same embryo in which a 6 nuclei wide patch of dorsal-most cells located at 50% EL were photoconverted at the blastoderm stage. (D) shows an embryo in which a 3 nuclei wide patch of dorsal-most cells located at 25% EL were photoconverted at the blastoderm stage. (E) shows an embryo in which a 3 nuclei wide by 6 nuclei long patch of dorsal-most cells located at roughly 2-10% EL were photoconverted at the blastoderm stage. In (A) and (C-E), embryos are oriented based on the AP/DV polarity of the egg with anterior to the left and dorsal to the top. In (A”), the surface of the egg is oriented to the bottom. In (B), the anterior of the germband is to the left. (A) is an average intensity projection of one egg hemisphere. (A’) is an average intensity projection of 6 μm to specifically show the dorsal epithelium. (B) is a maximum intensity projection of the whole germband. (C-E) are maximum intensity projections of one egg hemisphere. Abbreviations are: ventral ectoderm (VE) and dorsal ectoderm (DE). Scale bars are 50 μm.

At the end of GBE, all but the most posterior cells of the dorsal epithelium were squamous and *Tc-srp* seemed to be expressed in dorsal epithelium cells along nearly the full length of the germband (S4 Fig(I_2_)). However, this latter finding was difficult to confirm as most of the dorsal epithelium is lost during embryo fixation at this embryonic stage (presumably due to the fragility of the tissue). I also found *Tc-srp* to be expressed in several other domains, including in the presumptive endoderm (S4 Fig).

In *Drosophila, serpent* is also expressed in extraembryonic tissue (the amnioserosa) [42–45], and, therefore, *Tc-srp* may mark ‘true’ extraembryonic amnion. However, future work is required to confirm whether this putative amnion degenerates prior to hatching (as is required to be defined as extraembryonic). For simplicity, I will refer to this tissue as ‘amnion’ for the remainder of this text.

### A revised *Tribolium* amnion fatemap

To determine which blastoderm cells give rise to the amnion, I analysed 85 embryos in which the dorsal and dorsolateral blastoderm cells were labelled by NLStdEos photoconversion as described above. As I was unable to determine the precise DV position of the photoconverted cells at the blastoderm stage, I (1) examined embryos at the extended germband stage (when the mature amnion had formed), (2) determined the embryos in which photoconverted nuclei spanned the full DV width of the amnion (but were not observed in the embryonic tissue), and (3) checked the initial size of the photoconverted patch of nuclei (for a schematic of this approach, see S5 Fig).

I found that amnion cells arose from a very small domain of dorsal-most cells (that tapers from its anterior to posterior extent) and from a narrow strip of cells between the presumptive embryo and presumptive serosa (summarised in Fig 6 and S6 Fig). At 50% EL, only approximately the 6 most dorsal cells (approximately 8% of the circumference of the blastoderm) gave rise to all amnion cells stretching from one side of the thorax to the other (Fig 5(C), S1 Fig). Nearer to the posterior of the blastoderm (25% EL), even fewer cells gave rise to amnion (approximately 3 of the most dorsal cells; approximately 6% of the circumference; Fig 5(D), S2 Fig). The posterior limit of the amnion was difficult to define, as although some cells from approximately 5-10% EL appeared to become amnion (Fig5(E)), these cells condensed posteriorly towards the hindgut during germband retraction, and might have contributed to the hindgut tissue (S3 Fig, S7 Movie). I was unable to unambiguously determine the fate of these cells. At the anterior of the embryo, I found that a narrow strip of 1-2 cells between the presumptive embryo and presumptive serosa also gave rise to amnion (Fig 3(G-J)). While substantial additional work will be required to define a complete blastoderm fatemap for *Tribolium*, my findings clearly demonstrate that the ‘amnion’ domain is drastically smaller than previously proposed.

**Fig 6.**
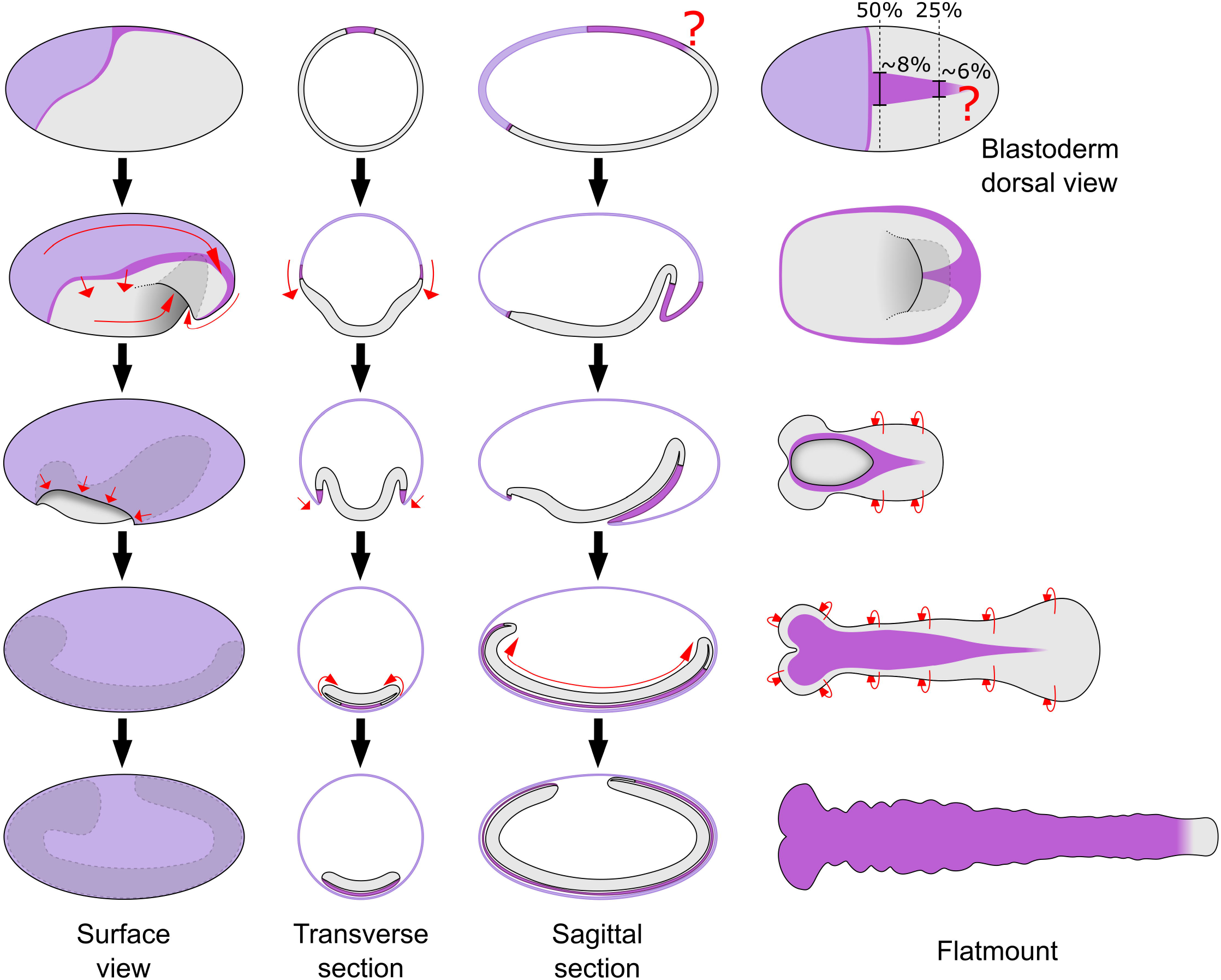
Schematics showing the revised *Tribolium* amnion fatemap and germband model. Schematics drawn as in Fig 1 to show the revised fatemap (top row; drawn directly from the numbers described in the text) and germband model based on the results of this manuscript. Note that the posterior amnion/embryo boundary is unclear. The schematics of the flatmounted germbands are drawn with the focus on the dorsal epithelium. See text for additional details, and S6 Fig for an extended figure with the classic and revised models side-by-side.–

## Discussion

In this article, I have shown that a majority of the cells currently thought to be extraembryonic amnion actually give rise to embryonic tissue. Movement of these cells from the dorsal side of the germband to the ventral side was visible in live cell tracking and differential cell labelling experiments. My results also indicate that the true amnion region differentiates progressively along the AP axis during GBE, as evidenced by differences in cell behaviour and the expression of the gene *Tc-srp*. Lastly, presumptive amnion cells predominantly originate from a small domain on the dorsal side of the blastoderm.

### A revised understanding of the short germ embryo

The revision to the *Tribolium* blastoderm fatemap that I describe is essentially a quantitative shift in our understanding of where cell fate boundaries lie along the DV axis. In the revised fatemap (Fig 6, S6 Fig), the proportion of the blastoderm that gives rise to the presumptive amnion is much smaller than previously thought. The presumptive amnion domain is, therefore, remarkably similar in size to the amnioserosa domain of the *Drosophila* blastoderm fatemap [18]. However, it is important to recognize that fatemaps such as those presented here show a static picture of a dynamic process. There is no evidence that the presumptive amnion is specified at the blastoderm stage in *Tribolium*. Instead, the progressive changes in cell shape and activation of *Tc-srp* expression in the dorsal epithelium of the germband suggest that the amnion is specified progressively along the AP axis during GBE. Progressive specification of DV cell fates during GBE fits with previous hypotheses [36,46], and analysis of how this process occurs represents an exciting avenue of future research (a possible mechanism for DV patterning during GBE is discussed in S1 Text).

In contrast to the fatemap revision, the observation that cells move from the dorsal half of the germband to the ventral half of the germband represents a qualitative shift in our understanding of development in short/intermediate germ insects. In the classic model of short/intermediate germ development, the germband was thought of as a more-or-less flat sheet of ectodermal cells (with mesoderm underneath) covered by the extraembryonic amnion. Because of this, the entire dorsal epithelium is routinely removed during embryo preparation, or not included in descriptions of gene expression patterns and embryonic phenotypes. Based on the new data presented here, it is obvious that we have been discarding or ignoring large parts of the embryo. Furthermore, the movement of cells from the dorsal epithelium into the ventral epithelium must be contributing to GBE, and is, therefore, a key aspect of the extension and overall development of the germband that has thus far been missed.

The revised model of the germband does present some technical challenges for future work on short/intermediate germ embryo. The flattened geometry of the germband makes it difficult to image both the dorsal and ventral epithelium using bright-field microscopy approaches. However, this problem can be overcome either by using fluorescence based techniques and confocal microscopy or by mechanical sectioning of the germband. Both approaches have been shown to work well in *Tribolium* (for examples see [13,47] and the results in this manuscript). In the rest of this article, I discuss why the revised fatemap and cell flow accord well with what we know about *Tribolium* development, and outline the implications of this discovery on our understanding of the evolution of insect development.

### The cellular and molecular causes of tissue flow unify the blastoderm and the germband

The revised model of the *Tribolium* germband reconciles the blastoderm and germband stages. The ectoderm of the germband is a continuous epithelium, which means that the movement of cells from the dorsal epithelium to the ventral epithelium occurs as a tissue level ‘flow’. Such dorsal-to-ventral tissue flow also occurs during embryo condensation in *Tribolium* [14], and I propose that the flow is caused by largely the same morphogenetic processes at both stages. The evidence for this hypothesis is summarised here, but for an extended discussion see S2 Text.

Three morphogenetic processes contribute to dorsal-to-ventral cell flow in *Tribolium*, and at least two of the three occur at both the blastoderm and germband stages. First, mediolateral cell intercalation occurs at both stages and causes tissue-wide convergence (along the DV axis) and extension (along the AP axis). This process requires two *Toll* genes that are expressed in rings around the entire blastoderm and germband epithelium [17]. Second, tissue specific cell shape changes occur at both stages such that ventral/lateral cells become columnar and dorsal/dorsolateral cells become thinner (during condensation (S7 Fig)) or squamous (during GBE). The tissue level effect of these changes is contraction of the ventral/lateral ectoderm and spreading of the dorsal tissue. The flattening of dorsal/dorsolateral cells is likely regulated by BMP signalling, as not only does BMP activity correlate with the cell shape changes (see S2 Text), but functional disruption of BMP signalling components leads to uniform cell shape changes along the DV axis [25,48]. A third major morphogenetic event is gastrulation of the mesoderm. This occurs along the ventral midline, and as gastrulation occurs, the ectoderm moves ventrally to seal the gap left in the epithelium [47]. At the stage when a complete germband has formed, gastrulation is complete along most of the embryo. However, current data suggests mesoderm gastrulation may be ongoing in the SAZ [47]. If true, the ongoing invagination would contribute to tissue flow in this region.

It is important to note that while each of the events described here is involved in the dorsal-to-ventral tissue flow, no single event is absolutely required for it. In the absence of cell intercalation, embryo condensation and thinning of dorsal/dorsolateral ectoderm still takes place, yielding abnormally wide and short germbands [17]. In the absence of tissue specific cell shape changes, condensation occurs in a more radially symmetrical manner yielding a tube-shaped germband that undergoes segment specification and convergent extension [25,48]. Finally, both condensation and GBE are only mildly affected in the absence of mesoderm specification [49]. This functional independence comes from each of the three processes being specified by different pathways (intercalation via segment specification, dorsal thinning via dorsal tissue specification, and gastrulation via ventral tissue specification). There may also be further, as yet undiscovered, morphogenetic events which also contribute to the dorsal-to-ventral tissue flow.

### Reconciling long and short germ development

I propose that the dorsal-to-ventral tissue flow occurring during embryo condensation and GBE in *Tribolium* is homologous to the dorsal-to-ventral tissue flow that occurs during gastrulation and GBE in *Drosophila* (Fig 1). This conclusion is based on the flow being driven by a conserved set of morphogenetic events.

As described above, tissue flow in *Tribolium* is caused by (1) mediolateral cell intercalation, (2) tissue specific cell shape changes along the DV axis, and (3) gastrulation at the ventral side of the embryo. As described below, equivalent processes are all observed in *Drosophila* as well.

In *Drosophila, Toll-*mediated mediolateral cell intercalation causes tissue-wide convergence (along the DV axis) and extension (along the AP axis) of the ectoderm during GBE [16]. As in *Tribolium*, the periodic expression of the *Toll* genes is regulated by the pair-rule genes. Conservation at the level of tissue identity, morphogenetic process, and molecular control strongly suggest *Toll*-mediated cell intercalation to be homologous.

Cell shape changes are harder to compare between *Drosophila* and *Tribolium*, because unlike in most insects, cellularisation in *Drosophila* leads to the direct formation of columnar cells [18,50]. However, tissue-specific cell shape changes along the DV axis do occur in *Drosophila* and are dependent on BMP signalling ([51,52]; for a detailed description see S3 Text). While the intracellular effectors of these cell shape changes are unknown, use of BMP signalling for dorsal patterning is homologous in *Drosophila* and *Tribolium*, and many dorsal cell specification genes are conserved between these two species [48].

Last, *Drosophila* mesoderm gastrulation also occurs along the ventral midline, and causes lateral/dorsolateral ectoderm to move ventrally [51]. Similar to the tissue specific cell shape changes described above, the intracellular effectors of *Tribolium* mesoderm gastrulation are unknown, but the upstream patterning events and the tissue specification genes are highly conserved [36,49]. Furthermore, mesoderm gastrulation at the ventral region of the embryo is widely observed within the insects, and is undoubtedly a homologous process in each species [53].

While I have focused on *Tribolium* and *Drosophila* here, evidence exists that the new findings in *Tribolium* may also apply to other short/intermediate germ insects. For example, in the intermediate germ bug *Oncopeltus fasciatus* (which forms a condensed, multi-layered germband with tissue topology similar to that of *Tribolium* [26]), the dorsal epithelium of the germband initially consists of a thick epithelium which progressively becomes squamous late during GBE [54]. These tissue-specific cell shape changes are likely the same as those occurring during *Tribolium* GBE. Furthermore, *Oncopeltus* pair-rule genes, Loto *Toll* genes and even segment polarity genes are expressed in rings around the entire germband prior to thinning of the dorsal epithelium [17,55,56]. The expression of these genes in the dorsal epithelium provides additional evidence that much of the *Oncopeltus* dorsal epithelium is made up of embryonic tissue. Future analyses of the molecular and morphogenetic drivers of GBE must analyse the entire germband, rather than focusing on the ventral half. In addition, further work will be needed to determine whether the new findings in *Tribolium* also apply to more basally branching insects such as crickets.

## Materials and Methods

*Tribolium* animal husbandry, egg collection, and RNA *in situ* hybridisation was performed as previously described [17]. The *Tc-srp* ortholog was previously described [57] and was cloned into pGEM-t (Promega Reference A1360) with primers TCCCGCTGCTTTGATCTAGT and TGCGATGACTGTGACGTGTA. The *Tc-cad* ortholog was as previously used [14].

The *H2B-ven* fusion was created by fusing the *D. melanogaster* histone *H2B* coding sequence (without the stop codon) from the published H2B-RFP [14] to the *venus* fluorescent protein [58] and cloning into the pSP64 Poly(A) (Promega Reference P1241) expression vector. The *NLS-tdEos* fusion was kindly provided by Matthias Pechmann. Additional details and both plasmids are available upon request to M. Pechmann or myself. Capped mRNA synthesis was performed as previously described [14]. *H2B-ven* capped mRNA was injected at 1 μg/μL, *NLS-tdEos* capped mRNA was injected at 2-3 μg/μL.

Embryo microinjection was performed as previously described [14], with the following changes. Up to 100 dechorionated embryos were mounted on a rectangular coverslip (24 mm by 50 mm) that rested on a microscope slide. Water was allowed to dry off the embryos before they were covered in Voltalef 10S halocarbon oil and injected as usual. The coverslip (still resting on the slide) was then placed in a petri-dish (92 mm) containing a base layer of 1% agarose (dissolved in water) and placed at 30-32°C until the embryos were at the appropriate stage for imaging. The coverslip was then removed from the slide, inverted (so that embryos were face down), and quickly but gently placed on a lumox dish (50 mm; Sarstedt Reference 94.6077.410) that was sitting upside down. The corners of the coverslip rested on the raised plastic lip of the dish such that the membrane and embryos were close to each other but not touching. To ensure lateral stability of the coverslip during the timelapse recording, approximately 5-10 μL of heptane glue (made by soaking parcel tape in heptane) was placed at each corner. Additional Voltalef 10S halocarbon oil was then added to fill any remaining space between the coverslip and the oxygen permeable membrane. This contraption was then stuck to a microscope slide (using double sided tape) for imaging on an upright microscope. This last step may be unnecessary depending on the microscope stage and orientation.

Live imaging was performed on an upright Zeiss SP8 confocal microscope equipped with Hybrid detectors in the Biocentre Imaging facility (University of Cologne). Image stacks of 15-50 focal planes with z-steps ranging from 2-10 μm were taken with a 10x/0.3N A dry objective or a 20×/0.7NA multi-immersion objective at intervals of 5-45 minutes. The temperature of the sample during imaging could not be carefully regulated, but was typically between 25-28 degrees. While this lack of temperature control is not ideal, it does not affect the findings presented in this manuscript.

Photoconversion of NLS-tdEos protein was performed by constantly scanning the region of interest for 20-30 seconds with the 405 wavelength laser at low power (5%). These settings were manually determined on the above microscope, and need to be determined independently on different systems. Photoconversions were performed during the final uniform blastoderm stage, as photoconversion prior to this resulted in substantial diffusion of the photoconverted protein during nuclei division. The positions of the different regions of the embryo (75% EL etc.) were determined by measuring the length of each embryo in the LASX software and selecting the appropriate region. Photoconversions were performed on all embryos on the coverslip before setting up the timelapse, which led to a 0.5-2 hour delay between performing the photoconversion and beginning the timelapse. As such, the positions of the photoconverted region at the first time point in the timelapses in this manuscript do not reflect the original region of photoconversion.

Imaging of fixed material was performed on an upright Zeiss SP8 confocal, an upright Zeiss SP5 confocal microscope and an inverted Zeiss SP5 confocal microscope. The *Drosophila gooseberry* expression patterns were kindly provided by Erik Clark and acquired as in [59]. Images and timelapses were analysed using FIJI [60] and Photoshop CS5. Manual cell tracking was performed on confocal hyperstacks with MTrackJ [61]. The figures were arranged and the schematics created using Inkscape.

## Acknowledgements

I am very grateful to S. Roth for his generous support, and for the extensive and stimulating discussions throughout this project. I am also thankful to E. Clark for in-depth discussions and extensive feedback on the manuscript. I thank M. Pechmann for providing the *NLS-tdEos* construct. In addition, I thank M. Akam, M. Pechmann and S. Roth for comments on the manuscript, and B. Schmitt for comments on the schematics.

---

**S1 Fig. Results of photoconversions at 50% egg length**. NLS-tdEos labelled extended germband stage *Tribolium* embryos in which a patch of blastoderm nuclei were photoconverted at 50% egg length (from the posterior pole) at different DV positions. The approximate DV position of the patch and the approximate DV width of the clone (in terms of nuclei number) are shown. The dorsal labelled embryo is shown from both sides to demonstrate the photoconverted nuclei cover the full DV extent of the amnion (arrows). Unconverted protein is shown in magenta, converted protein is shown in cyan. Images are maximum intensity projections of one egg hemisphere. All eggs are oriented with the anterior to the left and ventral to the bottom. Scale bars are 100 μm.

**S2 Fig. Results of photoconversions at 25% egg length**. NLS-tdEos labelled extended germband stage *Tribolium* embryos in which a patch of blastoderm nuclei were photoconverted at 25% egg length (from the posterior pole) at different DV positions. The approximate DV position of the patch and the approximate DV width of the clone (in terms of nuclei number) are shown. Unconverted protein is shown in magenta, converted protein is shown in cyan. Images are maximum intensity projections of one egg hemisphere. All eggs are oriented with the anterior to the left and ventral to the bottom. Scale bars are 100 μm.

**S3 Fig. Results of photoconversions near the posterior pole**. NLS-tdEos labelled extended germband stage *Tribolium* embryos in which a patch of blastoderm nuclei were photoconverted near the posterior pole at different DV positions. The approximate DV position of the patch and the approximate DV width of the clone (in terms of nuclei number) are shown. The second dorsally labelled embryo is shown at high magnification at two timepoints and with a transverse section (at the position of the dashed green line) to show the movement of tissue from the dorsal epithelium into the hindgut. Unconverted protein is shown in magenta, converted protein is shown in cyan. Images are maximum intensity projections of one egg hemisphere except for the bottom three embryos, which are shown as maximum intensity projects through the germband in order to better show the labelled nuclei. All eggs are oriented with the anterior to the left and ventral to the bottom except for the second timepoint of the second dorsal view, which is shown with the posterior of the germband to the left. Scale bars are 100 μm.

**S4 Fig. RNA expression of the *Tribolium* ortholog of the GATA factor *serpent***. (A-F) whole mount and (G-J) flatmount *Tribolium* embryos from the pre-blastoderm to the retracting germband stage stained for *Tc-srp* mRNA (red) and nuclei (DAPI, blue). (G_1_) and (G_2_) show the same embryo imaged from both sides. (H_1_) and (H_2_) show projections from the dorsal epithelium (H_1_) and the ventral epithelium (H_2_) of the same embryo. *Tc-srp* mRNA is maternally provided (A), and expression is ubiquitous until the late blastoderm stage (B-C) when expression clears from the blastoderm but persists in the yolk nuclei (scattered spots in (D-E)). During embryo condensation, *de novo* expression arises in a patch of blastoderm cells at the anterior medial region (arrowhead in F). This patch of *Tc-srp* expressing cells invaginates as part of the ventral furrow and becomes located beneath the ectoderm (arrowhead in G_1_-H_3_). This expression domain is likely homologous to the anterior ventral expression domain in *Drosophila* that marks the prohemocytes. During serosa window closure, expression appears in a ring of dorsal epithelium cells (G_1_). After serosa window closure, expression persists in the dorsal epithelium (H_1_) and (H_3_). Unlike *Drosophila*, there is no expression domain at the posterior of the blastoderm (E) or the early germband (G_2_). After germband elongation, a *de novo* expression domain appears at the posterior most point of the embryo (l_1_). Given the location of this domain at the base of the forming hindgut, this is likely the posterior endoderm primordium. Expression can also be seen in a patch of amnion that has remained attached to the germband (arrow in l_2_), but most of the rest of the amnion has been lost. Several other regions of expression can be seen, including in the presumptive fat body (the segmental domains running down the body), in presumptive hemocyte clusters (the two side-by-side domains in the anterior), and in an anterior domain that may mark the anterior endoderm primordium. Expression also persists in the yolk nuclei (visible in the remaining yolk fragments at the anterior of the germband in l_1_). All embryos are shown with anterior to the left. (E-E”) is oriented with the ventral to the bottom, (F-F”) is oriented as a ventral view. (H_3_-H_3_”) is an optical sagittal section at approximately the midline of the embryo in (H_2_). (A-F) are maximum intensity projections of one egg hemisphere. (G_1_-G_2_”), (l1-l1”) and (J-J”) are maximum intensity projections of flatmounted germbands. (l_2_-l_2_”) is a maximum intensity projection of part of the germband to better show the amnion/dorsal epithelium. Scale bars are 50 μm.

**S5 Fig. Schematics showing the photoconversion approach to determine the amnion fatemap**. A patch of nuclei (of known dimensions) was photoconverted at the blastoderm stage, then the same embryos were examined at the end of germband extension. In embryos where all photoconverted nuclei were located in the amnion and these nuclei spanned the entire DV width of the amnion (as shown here), the number of nuclei initially photoconverted was used to determine the DV width of the blastoderm domain giving rise to the amnion. Note that the precise number and distribution of nuclei shown here were arbitrarily chosen. Blue shows photoconverted nuclei, yellow shows the yolk. The serosa is omitted from the bottom panels.

**S6 Fig. Extended schematics showing the classic and revised *Tribolium* amnion fatemaps and germband models**. Schematics drawn as in Fig 1 to show the classic and revised fatemaps and germband models based on the results of this manuscript. The schematics of the flatmounted germbands are drawn with the focus on the dorsal epithelium. See text for additional details.

**S7 Fig. Tissue specific cell shape changes during *Tribolium* condensation**. Stills from timelapses of two *Tribolium* embryos transiently expressing GAP43YFP to label membranes. The second panel of each timepoint shows optical transverse sections at the position of the dashed line in the related panel. Ventral and lateral ectoderm becomes columnar, while dorsal ectoderm becomes flattened. The non-columnar cells at the bottom of the left hand embryo are likely the presumptive mesoderm. The first frame of the timelapses was defined as timepoint 0. Both embryos are oriented with the anterior to the left and ventral to the bottom. Abbreviations are: Dorsal (Dor), Lateral (Lat), Ventral (Ven) and Ectoderm (Ect). Scale bars are 100 μm.

**S1 Movie**. Confocal timelapse of a *Tribolium* embryo transiently expressing H2B-ven to mark nuclei. A maximum intensity projection of one egg hemisphere is shown. Anterior is to the left, the ventral side of the egg is to the bottom.

**S2 Movie**. Same timelapse as Supplementary Movie 1, but with nuclei of the dorsal epithelium tracked until they join the ventral epithelium. Nuclei that join the ventral epithelium are labelled magenta, nuclei that become located at the edge of the germband are labelled yellow, nuclei that remain in the dorsal epithelium are labelled cyan. Anterior is to the left, the ventral side of the egg is to the bottom. See Fig 2(A-C) for more details.

**S3 Movie**. Same timelapse as Supplementary Movie 1, but with a line of nuclei of the dorsal epithelium tracked. Nuclei that remain in the dorsal epithelium are labelled cyan. Anterior is to the left, the ventral side of the egg is to the bottom. See Fig 2(D-F) for more details.

**S4 Movie**. Confocal timelapse of a *Tribolium* embryo transiently expressing NLS-tdEos (magenta) with a line of blastoderm cells photoconverted (cyan). The brightness increases approximately halfway through the movie due to a manual increase in laser power at this point. A maximum intensity projection of one egg hemisphere is shown. Anterior is to the left, the ventral side of the egg is to the bottom.

**S5 Movie**. Confocal timelapse of a *Tribolium* embryo transiently expressing NLS-tdEos (magenta) with a patch of blastoderm cells photoconverted (cyan). (A) shows a maximum intensity projection of one egg hemisphere is shown. (B) shows an optical transverse section. (C) shows an average intensity projection of 10 optical transverse sections. Anterior is to the left, the ventral side of the egg is to the bottom. See Fig 3(A-F) for more details.

**S6 Movie**. Confocal timelapse of a *Tribolium* embryo expressing NLS-tdEos showing the SAZ during abdominal segment formation. Coloured points mark tracked nuclei. The embryo is oriented based on polarity of the visible region of the germband with posterior to the right and lateral to the top. Panels show maximum intensity projections of 15 μm to specifically show the germband. See Fig 4 for more details.

**S7 Movie**. Confocal timelapse of a *Tribolium* embryo transiently expressing NLS-tdEos (magenta) with a patch of blastoderm cells photoconverted (cyan). Towards the end of the timelapse, the cyan nuclei in the dorsal epithelium appear to condense posteriorly to the hindgut. A maximum intensity projection of one egg hemisphere is shown. Anterior is to the left, the ventral side of the egg is to the bottom.

**S1 Text. Extended discussion of topics from Discussion section ‘A revised understanding of the short germ embryo’**.

**S2 Text. Extended discussions of topics from Discussion section ‘Cellular and molecular causes of tissue flow’**.

**S3 Text. Extended discussions of topics from Discussion section ‘Reconciling long and short germ development’**.

